# Minimal immune cell subset differences in a cohort of close contacts of tuberculosis index cases

**DOI:** 10.1101/2025.08.04.668387

**Authors:** Sudhasini Panda, Catherine Cheng, Naomi Hillery, Donald G. Catanzaro, Nelly Ciobanu, Valeriu Crudu, Timothy Rodwell, Antonino Catanzaro, Julie G Burel, Bjoern Peters, Cecilia S. Lindestam Arlehamn

## Abstract

Understanding the perturbations in immune response across the spectrum of TB infection is still unclear. In this study, we followed a cohort of close contacts of pulmonary TB patients with serial QFT testing at 0, 3, 6, and 12 months, and stratified them into six subgroups: QFT-increasing (low/high), QFT converters (QFT- to QFT+), QFT+ stable, and QFT- individuals. Despite these distinct QFT trajectories, we observed minimal differences in immune cell frequencies, activation profiles, and T helper subset distributions among the QFT subgroups, suggesting limited immunological stratification based on QFT dynamics alone. Ex vivo immune phenotyping, including analysis of CD4, CD8, and NKT cell frequencies, memory T cell subsets, and activated T cells (HLA-DR⁺CD38⁺), failed to distinguish between QFT subgroups. Antigen-specific CD4 T cell responses assessed by the activation-induced marker (AIM) assay were elevated in QFT+ compared to QFT- individuals. These findings suggest that blood-based immune profiling may not capture subtle immunological transitions among QFT converters or individuals with increasing QFT responses. In contrast, individuals with active TB (ATB) showed clear immune perturbations. ATB patients at diagnosis exhibited significantly elevated frequencies of antigen-specific CD4 T cells, increased activated T cells, and higher frequencies of intermediate monocytes and NK cells compared to QFT+/QFT- contacts. Many of these immune features declined with treatment, indicating therapy-associated immune resolution. Additionally, shifts in T helper subsets and effector memory populations were observed over the course of treatment. These results suggest that while ex vivo immune profiling can robustly distinguish active TB from non-diseased states, it lacks the resolution to differentiate QFT subgroups based on QFT dynamics alone. This could reflect either immunological similarity among close contacts regardless of QFT status or limitations of blood-based phenotyping in detecting early or subclinical immune shifts. Our study highlights the challenge of immunologically distinguishing QFT-defined subgroups within close contacts using conventional ex vivo profiling approaches.

## INTRODUCTION

The course of *M. tuberculosis* (Mtb) infection is variable and heterogeneous in humans, driven by factors like host immunity, bacterial characteristics, and environmental conditions. According to the Delphi consensus framework, TB is a dynamic spectrum, moving beyond the traditional binary classification of latent and active disease (1). It encompasses a wide range of infection states, starting with individuals who remain uninfected or successfully clear Mtb shortly after exposure, without developing immunological evidence of infection. In individuals who become infected, TB can progress along a spectrum—from an early asymptomatic phase with a risk of progression, to subclinical TB where the bacteria are active but no symptoms are evident, and eventually to clinical TB (or active TB, ATB), characterized by noticeable symptoms and confirmed bacterial presence. The spectrum also accounts for individuals who have undergone treatment and achieved disease resolution (1). Immunological evidence for Mtb exposure and/or infection is defined as a positive immune response to Mtb antigens that can be measured either by the tuberculin skin test (TST) or by interferon-γ release assays (IGRA), which include Quantiferon TB Gold plus (QFT-Plus) and T-spot.TB (2,3). Both types of tests have drawbacks. TST may yield positive results in individuals with TB infection, BCG vaccination, or exposure to nontuberculous mycobacteria, whereas IGRAs mostly avoid these cross-reactions. None of the tests can distinguish between Mtb exposure, infection, and ATB disease (4,5), and they require a 6–8-week window post-exposure, limiting their ability to provide timely diagnosis and treatment. Neither TST nor IGRA can predict the subsequent development of ATB among high-risk contacts of pulmonary TB (6).

In the spectrum of TB, immune cells play a central role in determining infection outcomes and disease progression. Upon exposure to Mtb, the host immune system initiates a complex response involving all facets of the immune system. As the first response, innate immune cells such as macrophages, dendritic cells, and natural killer (NK) cells recognize and attempt to contain the infection through phagocytosis, cytokine release, and cytotoxic activity (7). Through co-evolution with humans, Mtb has developed mechanisms to evade the immune response, leading to a careful balance between bacterial containment and immune evasion. This delicate equilibrium is maintained mainly through a T cell-mediated response that controls bacterial replication and prevents disease progression. In ATB disease, immune response dysregulation often results in uncontrolled bacterial replication, tissue damage, and clinical symptoms. During Mtb infection and throughout treatment, the host immune profile undergoes dynamic changes reflecting bacterial burden and immune restoration. A study from our group had characterized a specific subset of CD4⁺ T cells expressing CCR6⁺CXCR3⁺CCR4⁻ (Th1*) in IGRA+ healthy individuals. These cells are notably expanded in those that are IGRA+ compared to healthy controls (8). ATB is typically marked by heightened inflammation, with elevated levels of pro-inflammatory cytokines and activated T cell subsets (9). As treatment progresses and bacterial load declines, these inflammatory signatures gradually resolve, giving way to a more regulated immune state (10,11). Tracking these shifts in immune markers offers valuable insights into disease progression, treatment response, and potential for relapse, highlighting the importance of immunological profiling in TB management and research (12–14).

Various studies have done immune profiling in ATB patients and IGRA+ individuals (15–17). Here, we determined the frequencies of immune cell subsets and the Mtb-specific T cell response in a cohort of individuals from the Republic of Moldova who are close contacts of ATB patients and thus at potentially high risk for developing ATB disease. Given the spectrum of TB infection, we focused on close contacts and stratified them based on longitudinal QFT responses to capture immune heterogeneity. These groups included: (i) persistently QFT- individuals, (ii) QFT+ at enrollment, (iii) QFT converters (i.e., from QFT- to QFT+), and (iv) individuals with fluctuating or varying QFT magnitudes. This stratification enabled us to examine shifts in immune cell phenotypes and Mtb-specific T cell responses and compare them to ATB patients, both before and after completion of anti-TB therapy. The most drastic differences were found in the ATB cohort, with changes in innate and adaptive immune cell subsets frequencies and a reduction in the magnitude of antigen-specific immune responses with anti-TB treatment. In contrast, minimal differences were found between the different cohorts of close contacts stratified by their QFT results.

## RESULTS

### A cohort of individuals with active TB and close contacts of a TB index case

In this study, we recruited patients with ATB at the time of diagnosis (n=40) and mid/end of treatment along with a group of close contacts of patients with ATB, with follow-up samples at 3 months, 6 months, and 12 months. This longitudinal cohort of close contacts allows for an in-depth investigation of the immune response against Mtb over time, as well as capture individuals across a spectrum of TB infection stages and outcomes. Close contacts of pulmonary ATB patients were monitored longitudinally with QFTplus testing at 3-, 6-, and 12-months post-identification of the case of ATB. Analysis of their QFT trajectories showed heterogeneous patterns of QFT response (Figure 1a). Some individuals showed an increased magnitude of QFT response over time, and a subset of participants converted from QFT- to QFT+ during the follow-up period. Additionally, others maintained consistent positive or negative QFT responses. To better capture this variability and facilitate downstream immunological analyses, we subdivided these individuals into six distinct groups based on their QFT: (I) QFT Increasing (n=57, Figure 1b), which were further divided into (Ia) QFT low and (Ib) QFT high, (II) Converters (n=41, Figure 1c) subdivided into (IIa) QFT- vs. (IIb) QFT+, (III) QFT+ Stable (n=42, Figure 1d), (IV) QFT- Stable (or QFT-) (n=50, Figure 1e), as described in detail in the Methods section. The latter two groups QFT+ stable and QFT- were both represented by one timepoint per participant. PBMC samples collected from this cohort of individuals had cell viability issues. To be included in downstream immune analysis, a cut-off of 50% viability after thawing of the PBMC samples was used. This limited the ability to investigate longitudinal changes for each close contact, but allowed cross-sectional comparisons between close contact cohorts and the ATB cohort at diagnosis versus treatment.

**Figure 1:**
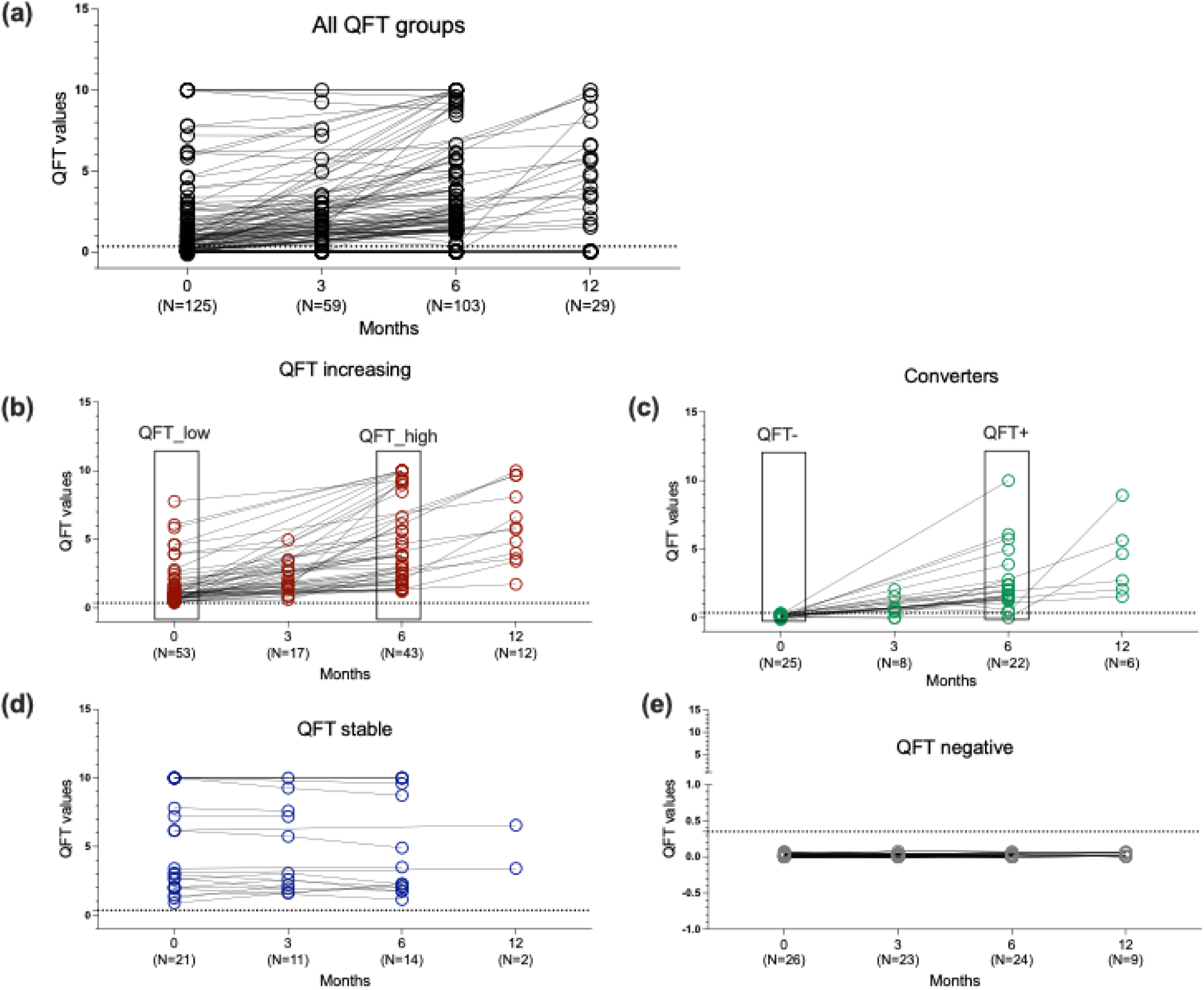
QFT values in close contacts of TB patients. (a) QFT values of all the individuals at 0, 3, 6, and 12 months. (b-e) QFT values of QFT increasing, converters, QFT+ stable, and QFT- stable individuals, respectively, at 0, 3, 6, and 12 months. (b) QFT increasing were also divided into QFT low (month 0) and QFT high (month 6). (c) Converters were also divided into QFT- (month 0) and QFT+ (month 6). (d) shows QFT stable whose QFT values remained stable over the time and (e) QFT- individuals whose QFT values remained negative over the time. A QFT value less than 0.35 IU/ml was considered QFT-. This is indicated by a dotted line in each graph.

### Minimal differences in T cell subsets across QFT subgroups and ATB

We determined ex vivo differences in T cell subset frequencies in the different above-mentioned cohorts. First, we compared the differences between combined QFT+ groups with QFT- and ATB individuals, followed by comparisons among QFT subgroups stratified by QFT response magnitude as mentioned above. We found a significant reduction in CD4+ T cells in the ATB group compared to the QFT+ and QFT- groups, whereas the CD8+ T cell levels showed no significant difference across the cohorts (figure 2a, b). We also looked at the frequency of NKT cells, which remained comparable between ATB and QFT+/- groups but showed an increase during treatment for ATB (figure 2c). When we examined individuals stratified by QFT results, we observed comparable frequencies of CD4⁺, CD8⁺, and NKT cells across subgroups (Figures 2d–f). This indicates that peripheral frequencies of these T cell subsets do not vary based on QFT results alone and are insufficient to stratify this heterogeneity.

**Figure 2:**
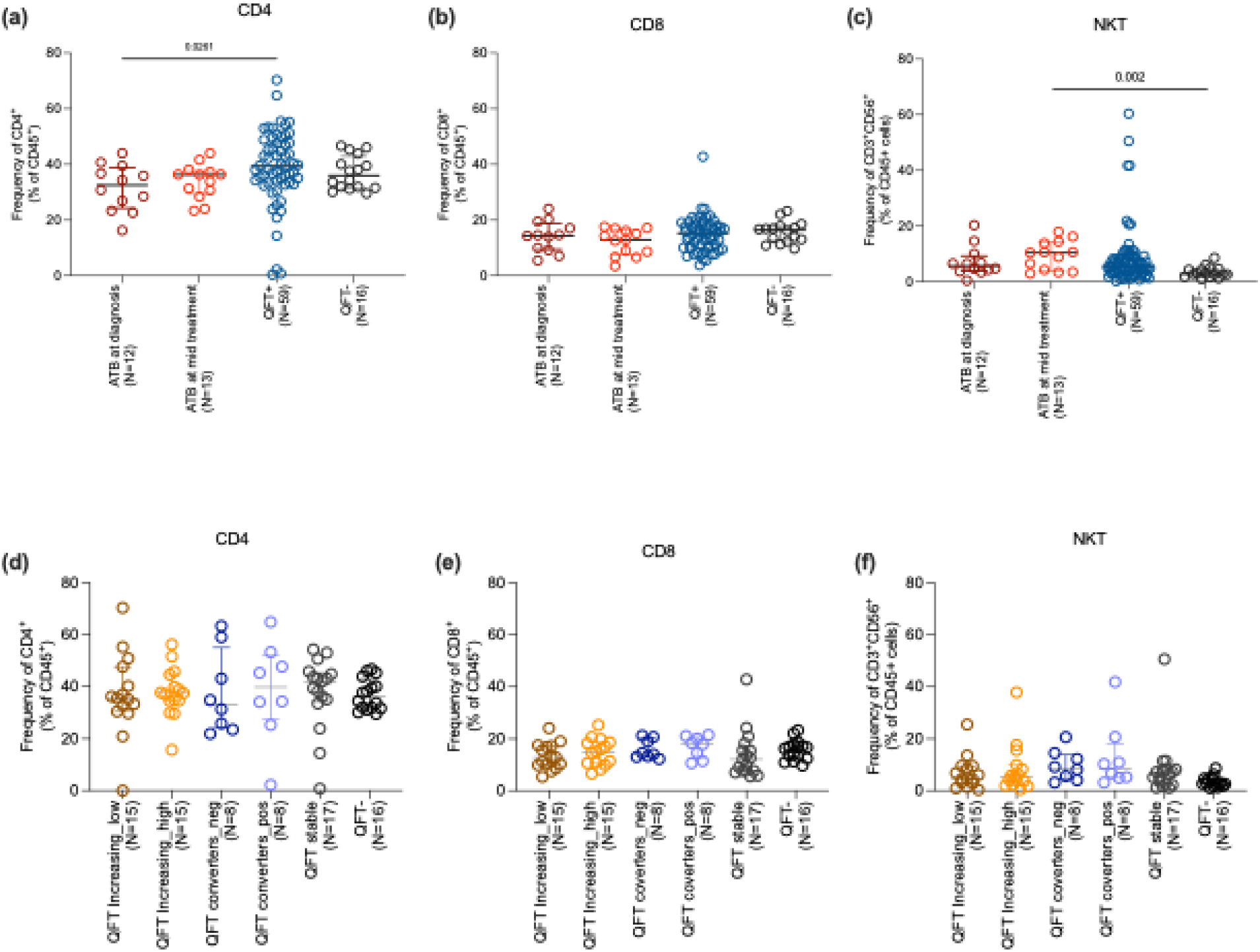
Ex vivo frequency of CD4, CD8 and NKT cells. (a,b and c) shows the frequency of CD4, CD8 and NKT cells respectively in ATB (at diagnosis and mid treatment), and in combined QFT+ and QFT- individuals. (c-d) shows the frequency of CD4, CD8 and NKT cells respectively in QFT stratified subgroups consisting of 6 groups: QFT increasing ((i)low and (ii)high), QFT converters ((iii)QFT- to (iv)QFT+), (v)QFT stable and (vi)QFT-, as defined in Figure 1. Each dot represents an individual participant, with the median and interquartile range indicated. Statistical comparisons were performed using the Kruskal-Wallis test followed by Dunn’s multiple comparison test. For comparison between two groups, the Mann-Whitney U test was used. p<0.05 was considered significant.

The overall frequency of CD4 memory subsets was also assessed using CD45RA and CCR7 markers. No significant differences were observed between ATB and QFT+/- cohorts (figure 3a-d), and among QFT subgroups (figure 3e-h).

**Figure 3:**
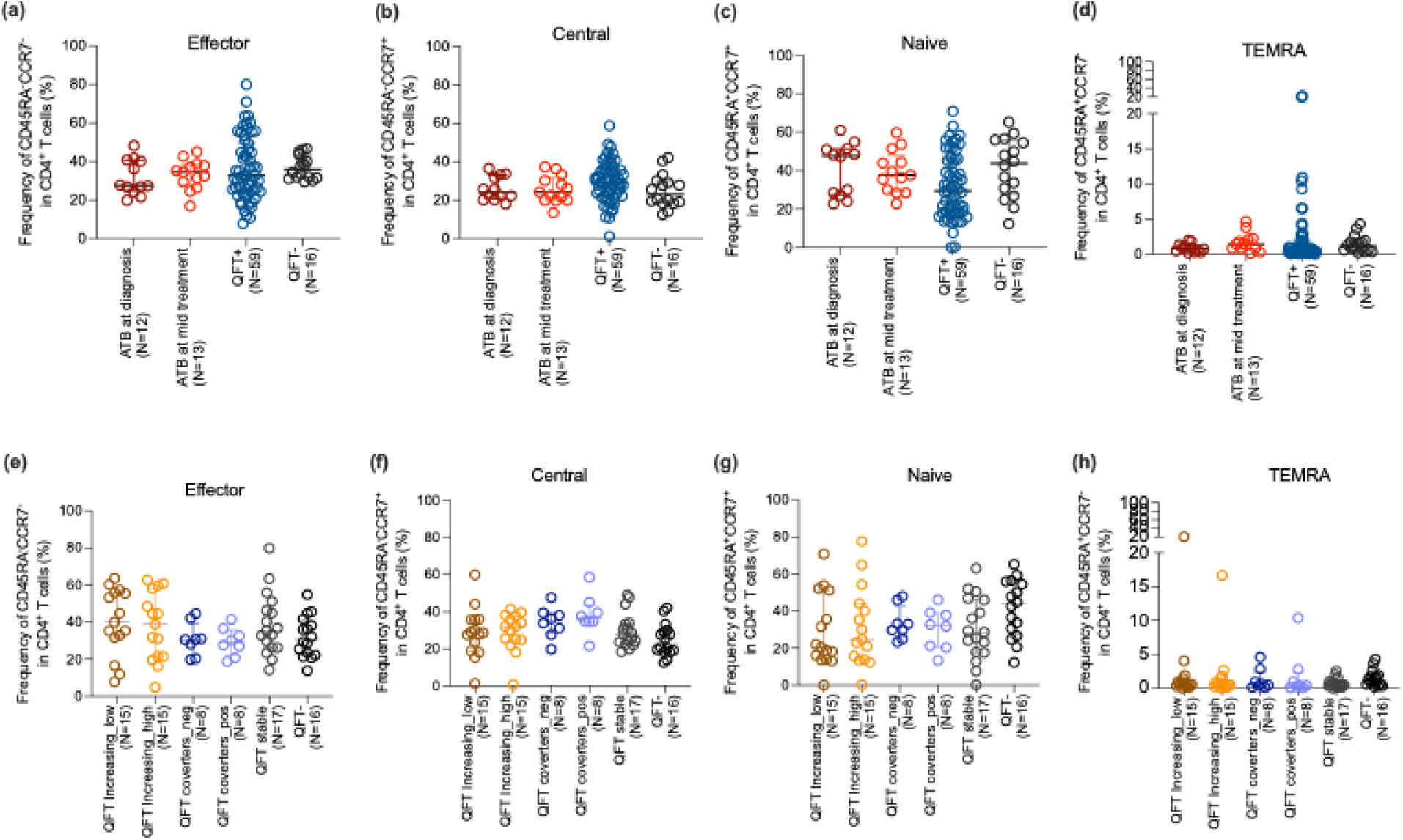
Frequency of memory CD4 T cell subsets. CD4 T cell subsets were defined according to surface expression of CD45RA and CCR7 with (i) CCR7-CD45RA- (effector memory), (ii) CCR7+CD45RA- (central memory), (iii) CCR7+CD45RA+ (naïve), and (iv) CCR7-CD45RA+ (T_EMRA_) T cells. (a-d) shows the frequency of each memory subsets in ATB (at diagnosis and mid treatment), and in combined QFT+ and QFT- individuals. (e-h) shows the frequency of each memory subsets in QFT stratified subgroups consisting of 6 groups: QFT increasing ((i)low and (ii)high), QFT converters ((iii)QFT- to (iv)QFT+), (v)QFT stable and (vi)QFT-, as defined in Figure 1. Each dot represents an individual participant, with the median and interquartile range indicated. Statistical comparisons were performed using the Kruskal-Wallis test followed by Dunn’s multiple comparison test. For comparison between two groups, the Mann-Whitney U test was performed. p<0.05 were considered significant.

We also assessed the frequency of recently activated CD4+ and CD8+ T cells using the markers HLA-DR and CD38 and found a higher frequency of HLA-DR+ CD38+ CD4 and CD8 T cells in the ATB group at diagnosis (figure 4a, b). These frequencies decreased with effective treatment, suggesting that the elevated activation levels were associated with the active infection state in ATB individuals. This observation matches previous studies, which have shown that the simultaneous expression of HLA-DR and CD38 on CD8+ T cells is particularly indicative of recent immune activation, such as during acute infections or early stages of treatment response in infections(22). No significant differences in the recent activation status of CD4+ or CD8+ T cells were observed in QFT+ or QFT- individuals. In the QFT subgroups, no difference was observed in the frequency of activated CD4 and CD8 T cells (figure 4c, d). This highlights that the heightened T cell activation is specifically related to the active disease process in ATB.

**Figure 4:**
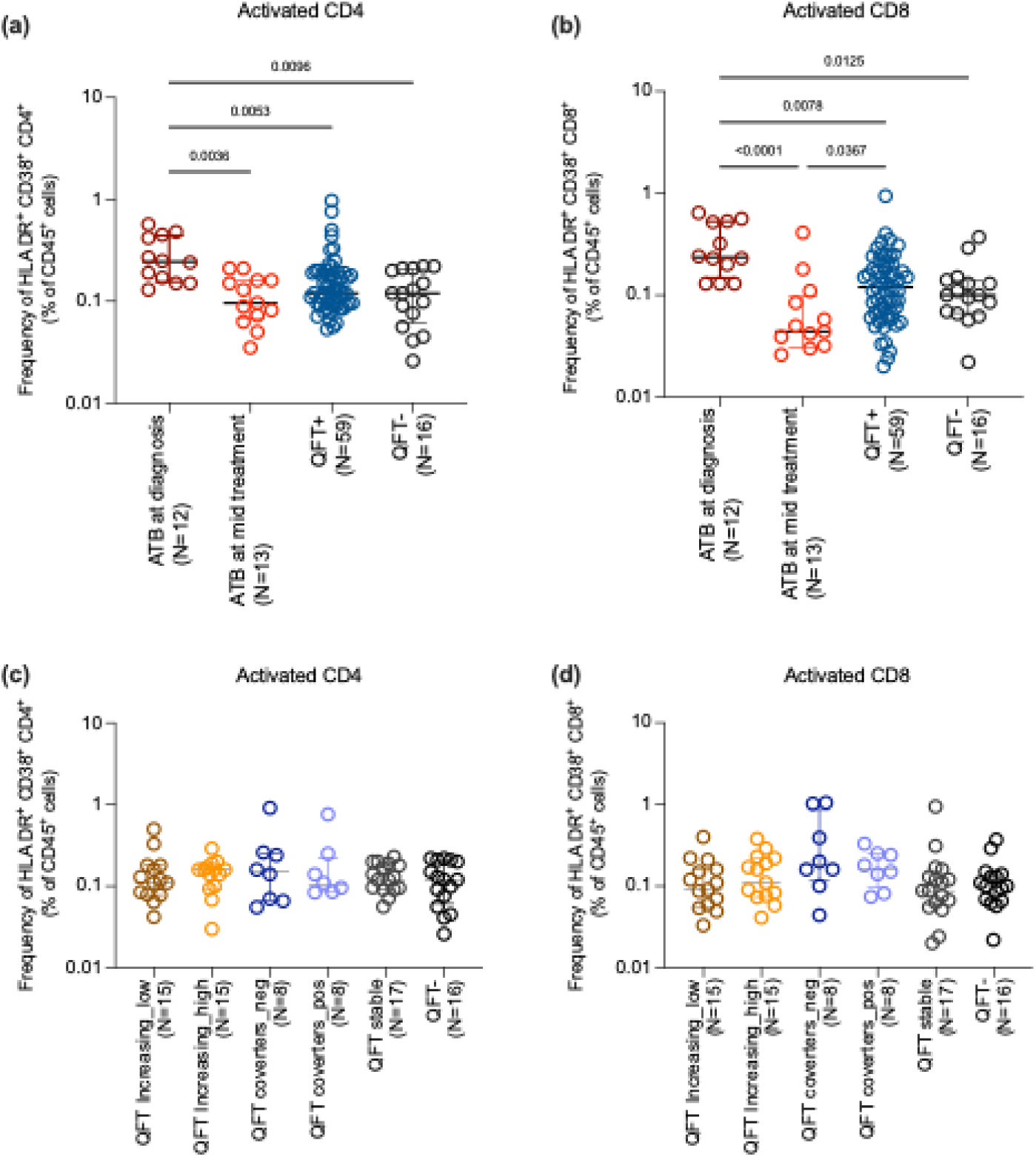
Frequency of activated CD4 and CD8 T cells: Activated CD4 T cells were defined as CD4+HLADR+CD38+ and activated CD8 T cells as CD8+HLADR+CD38+. (a-b) shows the frequency of activated CD4 and CD8 T cells in ATB (at diagnosis and mid-treatment), and combined QFT+ and QFT- individuals. (c-d) shows the frequency of activated CD4 and CD8 T cells in QFT stratified subgroups consisting of 6 groups: QFT increasing ((i)low and (ii)high), QFT converters ((iii)QFT- to (iv)QFT+), (v)QFT stable and (vi)QFT- as defined in Figure 1. Each dot represents an individual participant, with the median and interquartile range indicated. Statistical comparisons were performed using the Kruskal-Wallis test followed by Dunn’s multiple comparison test. For comparison between two groups, the Mann-Whitney U test was performed. p<0.05 was considered significant.

Together, our results indicated a higher activated state at diagnosis compared to mid-treatment in ATB, with increased frequencies of HLADR/CD38 CD4 and CD8 T cells but no differences among QFT subgroups. In particular, those who converted their QFT during the follow-ups did not look more similar to ATB diagnosis as we would expect. These findings imply that while ex vivo profiling effectively distinguishes ATB from non-diseased individuals (QFT+/-), it lacks the resolution to differentiate between QFT+ subgroups based on their QFT trajectories or risk of progression.

### Distinct Th Subset Dynamics specific to ATB Highlight Immune Modulation and Treatment Effects

Specific Th subsets have been described to be involved in Mtb-specific responses. To determine whether any of these participant cohorts had alterations in their Th subsets, we classified Th subsets based on their chemokine markers, CCR6, CXCR3, and CCR4. Th1* cells exhibit a CCR6+CXCR3+CCR4- phenotype and share lineage-specific characteristics with conventional Th1 (CCR6-CXCR3+CCR4-) and Th17 (CCR6+CXCR3-CCR4+) cells, whereas Th2 cells were defined as CCR6-CXCR3-CCR4+. There was no difference in the frequency of Th1* cells between ATB at diagnosis and QFT+/- groups (figure 5a), in contrast to what has been described previously (8). However, the frequency of Th1 cells was notably lower in ATB cases compared to QFT+ and QFT- individuals. (figure 5a, p-values are listed in supplementary table 1) There was also a decline in the frequency of Th1* and Th1 subsets during treatment. For the other Th subsets, there was a significantly higher frequency of Th17 cells in the ATB group compared to both QFT+ and QFT- groups. Th17 cells are associated with pro-inflammatory responses and are known to play a role in the pathogenesis of TB (23–25). The frequency of Th2 cells was also increased during treatment in comparison to QFT+ and QFT- groups. Interestingly, all the CXCR3+ subsets, irrespective of CCR4 and CCR6 expression, declined between diagnosis and mid-treatment ATB cohorts. There were no significant differences in the frequency of the different Th subsets between QFT- and QFT+ individuals (figure 5a). To investigate the balance of Th subsets in these participants in more detail, the ratio of Th1/Th2 was calculated and found to be lower in ATB compared to QFT+ and QFT- individuals (figure 5b), reflecting an altered immune balance characteristic of tuberculosis infection (26). For the QFT subgroups, similar to other subsets mentioned above, we did not find any significant differences in these Th subsets based on chemokine receptors (figure 5c-6e). Overall, the frequency of CXCR3+CCR6+ cells was lower than that of other single-positive or double-negative cells. This suggested that this blood-based ex vivo phenotyping could not distinguish these subgroups. The similarity between QFT subgroups may also stem from the fact that all participants were close contacts of TB patients, and thus exposed to Mtb with the potential for immune involvement, regardless of their QFT result.

**Figure 5:**
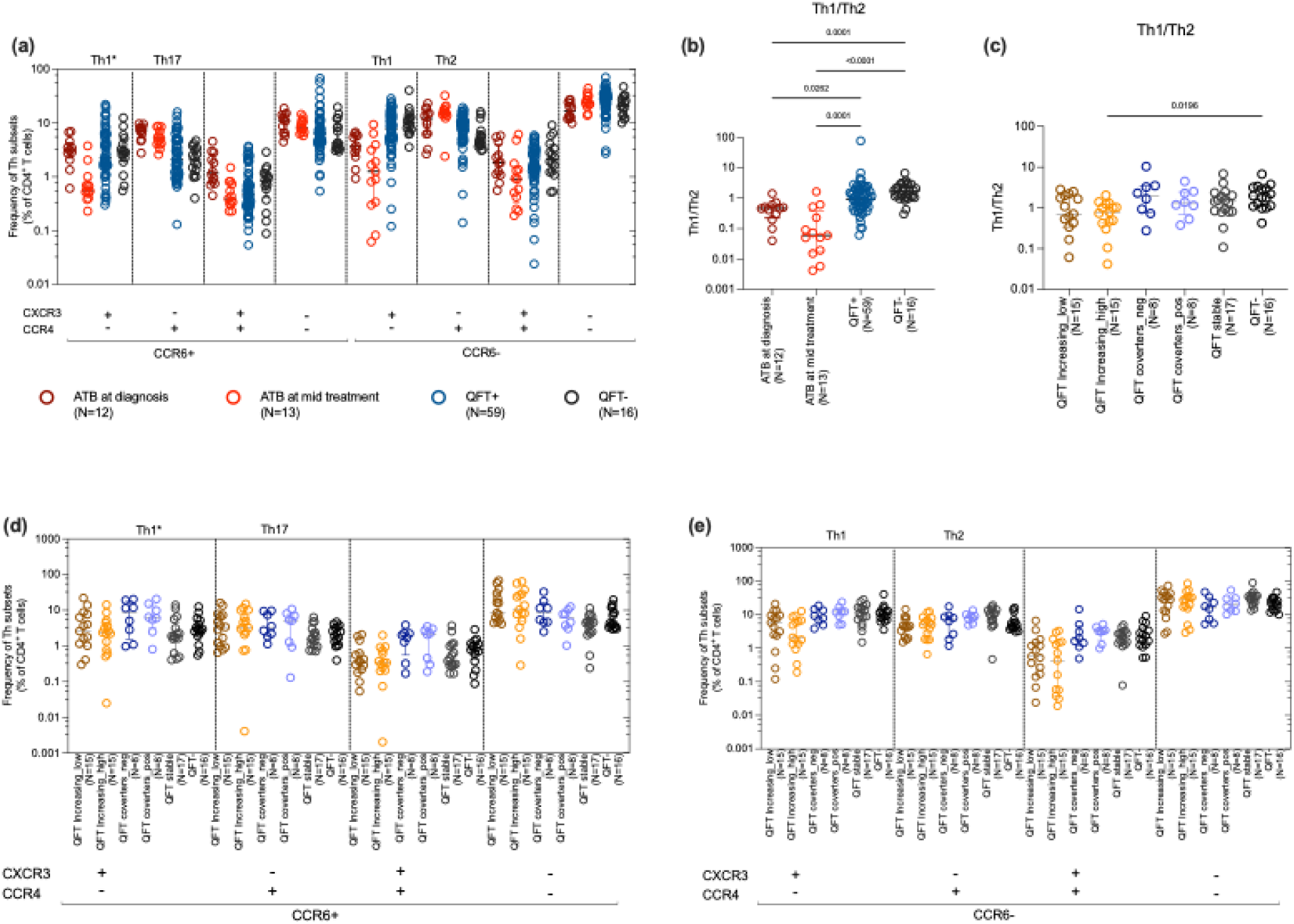
Ex vivo frequency of different T helper subsets study groups based on chemokine markers, CCR6, CXCR3 and CCR4. All subsets were gated from total CD4 T cells. Th1* was defined as CCR6+CXCR3+CCR4-. Th17 was defined as CCR6+CXCR3-CCR4+. Th1 was defined as CCR6-CXCR3+CCR4- and Th2 was defined as CCR6-CXCR3-CCR4+. (a) shows the frequency of all the Th based on expression of CXCR3, CCR6, and CCR4 in ATB (at diagnosis and mid-treatment), and in combined QFT+ and QFT- individuals. (b) shows the ratio of Th1 and Th2. (c) shows the ratio of Th1/Th2 in the same subsets in QFT stratified subgroups consisting of 6 groups: QFT increasing ((i)low and (ii)high), QFT converters ((iii)QFT- to (iv)QFT+), (v)QFT stable and (vi)QFT-, as defined in Figure 1). (d) and (e) shows the frequencies of the same subsets in QFT stratified subgroups Panel (d) includes all the CCR6+ subsets and (e) includes CCR6-subsets. Each dot represents an individual participant, with the median and interquartile range indicated. Statistical comparisons were performed using the Kruskal-Wallis test followed by Dunn’s multiple comparison test. For comparison between two groups, the Mann-Whitney U test was used. p<0.05 was considered significant.

**Figure 6:**
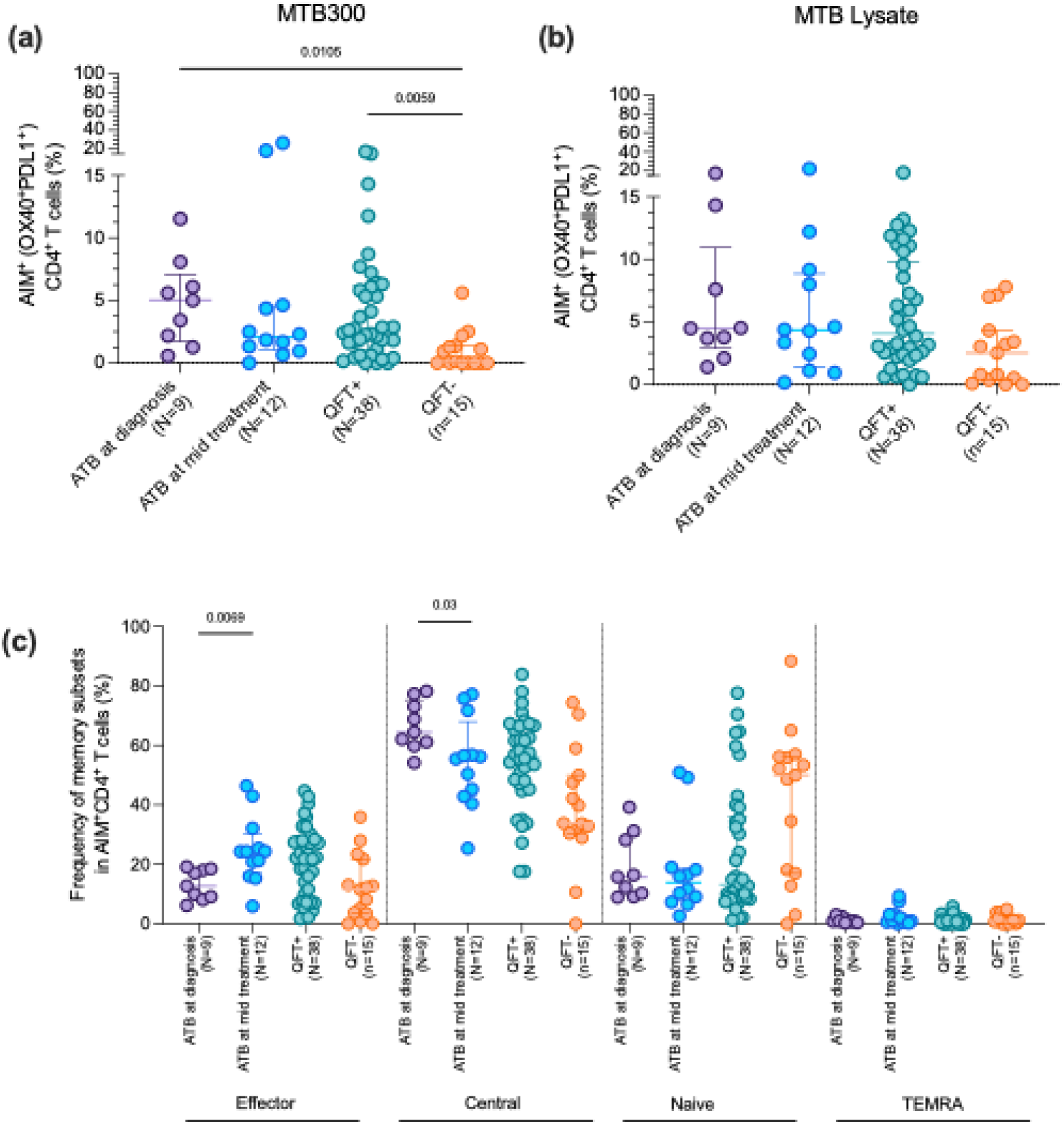
Frequency and phenotype of AIM+ CD4 T cells in ATB, QFT+, and QFT- individuals. AIM+ cells were selected based on the upregulation of activation-induced markers, namely OX40 and PDL1 on CD4 T cells. (a) and (b) shows the frequency of AIM+ cells after stimulation with MTB300 and MTB lysate respectively. (c) shows the frequency of memory subsets within AIM+ CD4 T cells. Each dot represents an individual participant, with the median and interquartile range indicated. Statistical comparisons were performed using the Kruskal-Wallis test followed by Dunn’s multiple comparison test. For comparison between two groups, the Mann-Whitney U test was performed. p<0.05 was considered significant.

Increased antigen-specific CD4 T cell response in participants with ATB

We have previously described the antigen-specific IFNγ reactivity against MTB300 (a peptide pool with 300 Mtb-derived T cell epitopes(18) and ATB116 (a peptide pool consisting of peptides that elicited IFNγ response specifically in individuals with ATB) between participants from this cohort with ATB, and QFT+ and QFT- close contacts(19). We found an increased IFNγ response in ATB individuals at diagnosis and mid-treatment compared to both QFT+ and QFT- controls after stimulation with the active specific peptide pool, ATB116(19). Here, we evaluated the antigen-specific response using the activation-induced marker (AIM) assay. This method identifies antigen-specific CD4 T cells by detecting the upregulation of surface markers following antigen stimulation, irrespective of which cytokine they produce. Specifically, we measured the expression of OX40 and PD-L1 after stimulation with MTB300 and Mtb whole cell lysate to allow capturing donor-unrestricted T cell responses as well. Previous studies have demonstrated that OX40+PD-L1+ cells are consistently induced at a higher frequency in QFT+ individuals compared to QFT− individuals after Mtb-specific stimulation and are absent in unstimulated samples (20,21). Due to limited sample availability for this analysis, the close contacts were divided into QFT+ vs. QFT- (38 QFT+ and 15 QFT-) rather than the 6 subgroups. As expected, we observed a higher frequency of AIM+ cells in ATB individuals and QFT+ compared to QFT− close contacts (p value = 0.01 and 0.005, respectively, Figure 6a). While the median frequency of AIM+ cells was lower in individuals with ATB mid-treatment compared to diagnosis, the difference was not significant (p=0.34). In addition, no significant differences were noted in the antigen-specific response after stimulation with Mtb lysate, which more broadly activates T cells, including non-conventional T cells (Figure 6b). These results demonstrate that the AIM assay can distinguish ATB and QFT+ individuals from QFT− individuals responding to MTB300 but not Mtb lysate.

After identifying significant differences in AIM+ cells, we assessed memory subset composition within these cells using CD45RA and CCR7 to define naïve (CD45RA⁺CCR7⁺), central memory (CD45RA⁻CCR7⁺), effector memory (CD45RA⁻CCR7⁻), and TEMRA (CD45RA⁺CCR7⁻) populations. Central memory cells were the most abundant subset across all cohorts, followed by effector memory cells (Figure 6c). Mid-treatment ATB patients showed a notable increase in effector memory AIM⁺ (p=0.006) cells alongside a significant decline in central memory subsets (p=0.03) compared to ATB at diagnosis, indicating that anti-TB therapy alters the composition of antigen-specific memory T cell responses. QFT+ individuals showed higher frequencies of both effector and central memory cells than QFT- individuals, while no significant differences were observed in the TEMRA subset (figure 6c). We were unable to assess differences in AIM+ cells across the QFT-stratified subgroups due to an insufficient number of data points for meaningful comparison. This was because of insufficient cell counts for 24 hour stimulation to the cells with MTB300 and Mtb lysate for the assay.

### Increased Intermediate Monocytes and Elevated NK Cells in Active TB, with Treatment-Associated Reduction

Finally, we analyzed the frequency of the most frequent non-T cell immune cell subsets in human blood across the cohorts, namely classical, intermediate, and nonclassical monocytes, and natural killer cells. We did not observe significant changes in CD56+NK cell frequencies between ATB at diagnosis and QFT+/- groups. However, we observed a higher frequency of NK cells in ATB at mid-treatment compared to QFT+ (p value = 0.03) (figure 7a). In terms of monocyte populations, while there was no significant difference in total monocyte frequencies across the cohorts, a higher frequency of intermediate monocytes was found in ATB individuals at diagnosis compared to mid-treatment (p value=0.001) (figure 7b). Intermediate monocytes have a unique phenotype and function, distinct from classical and non-classical monocytes, that are known to play a role in inflammatory responses (27). The frequency of intermediate monocytes in ATB patients decreased with treatment (figure 7b), indicating a potential normalization of immune activation as the infection is controlled, and in line with our previous single-cell profiling study of circulating monocytes in ATB and IGRA+(28). Importantly, the levels of intermediate monocytes were comparable between QFT+ and QFT- cohorts, corroborating the findings by Sampath et al., and suggesting that intermediate monocytes might serve as a specific marker for active TB disease (29). We observed comparable frequencies of NK and monocyte populations in the different QFT subgroups (figure 7d-7e).

**Figure 7:**
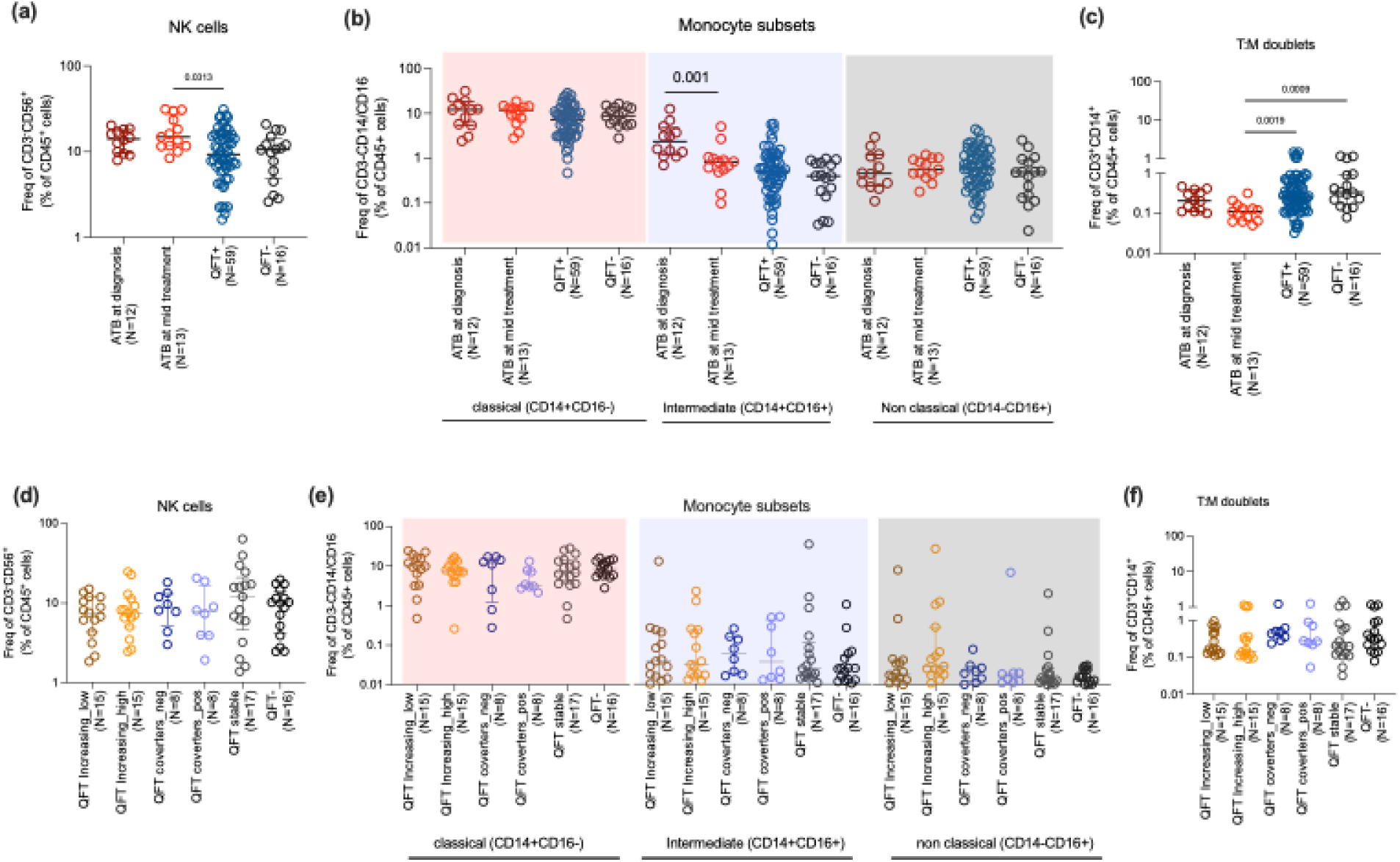
Frequency and phenotypic distribution of innate cells, NK cells, monocyte subsets and T:M doublets. (a) shows the frequency of NK cells (CD3-CD14-CD19-CD56+) in total leukocytes. (b) shows the frequency of different subsets of monocytes within CD3-CD19-CD56-cells: classical (CD14+CD16-), intermediate (CD14+CD16+) and non classical (CD14-CD16+). (c) shows the frequency of T:M doublets in ATB (at diagnosis and mid treatment), and in combined QFT+ and QFT- individuals (d-f) shows the frequencies of the same subsets in QFT stratified subgroups consisting of 6 groups: QFT increasing ((i)low and (ii)high), QFT converters ((iii)QFT- to (iv)QFT+), (v)QFT stable and (vi)QFT-, as defined in Figure 1. Each dot represents an individual participant, with the median and interquartile range indicated. Statistical comparisons were performed using the Kruskal-Wallis test followed by Dunn’s multiple comparison test. For comparison between two groups, the Mann-Whitney U test was used. p<0.05 was considered significant.

We also investigated the frequency of T cell-monocyte doublets (or T:M doublets). T:M doublets, pairing one T cell and one monocyte, are identified as CD3+CD14+ events within the singlet gate of human PBMC gated by flow cytometry (30). Their frequencies are typically increased following immune perturbations such as disease or vaccination and are enriched for recently activated T cells (31). No significant differences in T:M doublet frequencies were observed between ATB at diagnosis and the QFT+/- group (figure 7c). However, we found a significant decrease in T:M doublets at 2 months post-treatment compared to QFT+ and QFT- (p value= 0.001 and 0.0009 respectively, figure 7c), as previously observed (30). This reduction suggests that the presence of T:M complexes is associated with active infection and decreases as the infection is controlled through treatment. Similar to our other immune cell type measurements, we did not find any significant difference in T:M doublet frequency between the different QFT subgroups (figure 7f).

## DISCUSSION

This study aimed to characterize the immune response and immune cell subset frequencies across the spectrum of Mtb infection by examining close contacts of TB patients. We focused on close contacts of TB patients, incorporating QFT+ individuals stratified into subgroups based on longitudinal QFT dynamics to capture the immunological heterogeneity within TB infection. In addition to these groups, we also included pulmonary TB patients both at diagnosis and mid-treatment to examine treatment-associated immunological changes and to compare their immune profiles with those of QFT+ and QFT- individuals. Our findings contribute to the growing body of evidence on the heterogeneity of immune responses in TB and monitoring treatment efficacy. The analysis was performed in two stages: first by comparing broader categories, active TB (diagnosis and mid-treatment), QFT+, and QFT- individuals, and subsequently by examining immune parameters within the stratified QFT subgroups.

Our previous work demonstrated an enhanced antigen-specific CD4 T cell response in individuals with active tuberculosis (ATB) compared to QFT+ and QFT− controls, in response to stimulation with the MTB300 peptide pools (19). In this study, to capture as much of the antigen-specific immune response as possible, irrespective of cytokines produced, we utilized the Activation-Induced Marker (AIM) assay, focusing on OX40 and PD-L1 as markers of antigen-specific CD4 T cell activation (32). Previously, this assay has been used to detect antigen-specific and vaccine-specific CD4 T cell responses (33–35). As expected, a significantly higher frequency of OX40+PD-L1+ CD4 T cells in response to MTB300 stimulation was found in individuals with ATB and QFT+ individuals compared to QFT− controls. These results align with prior research, which emphasizes heightened immune activation in infection and diseased states, likely driven by the increased antigenic burden characteristic of ongoing infection (20). Notably, the frequency of AIM+ CD4 T cells is reduced during TB treatment, which suggests a response to the declining antigenic stimulation and immune activation as the infection resolves. These dynamics highlight the potential of the AIM assay not only to differentiate between active TB disease and latent TB infection (LTBI) but also to serve as a valuable tool for monitoring treatment response. Several studies have utilized different combinations of AIM markers, such as OX40, CD25, CD69, and CD40L, to identify Mtb-specific CD4⁺ T cells in a cytokine-independent manner, both in HIV-uninfected and HIV-infected individuals with active TB or latent infection, particularly in high TB burden settings (36,37).

There were distinct differences in AIM+ memory T cell subset frequencies during TB infection and treatment. AIM+ T cells were primarily central memory T cells, which showed a decrease during anti-TB treatment and a corresponding increase in effector memory cells. This phenotypic shift likely reflects the resolution of infection and a transition from sustained antigenic stimulation to a more immediate response phenotype. QFT+ individuals exhibited higher frequencies of both central and effector memory cells compared to QFT− controls, which could reflect the higher antigenic burden in these individuals. These findings highlight the role of memory T cell dynamics in immune status and disease progression. A previous study found that Mtb infection induces a population of functional, antigen-specific CD4+ stem cell memory T (T_SCM_) cells in humans. These T_SCM_ cells show long-lived memory characteristics, the ability to self-renew, and can give rise to more differentiated T cell subsets, suggesting they may play an important role in long-term immunity and protection against TB (38). We were unable to assess differences in AIM+ cells across the QFT-stratified subgroups due to an insufficient number of data points for meaningful comparison.

We initially hypothesized that individuals with varying QFT values might exhibit distinct immune cell compositions. However, we did not observe significant differences in ex vivo immune cell frequencies among the QFT-stratified subgroups. This suggests either that these subgroups do not differ substantially in their immune profiles or that conventional ex vivo phenotyping lacks the sensitivity to resolve such differences. In contrast, clear differences were evident when comparing immune cell frequencies across broader clinical categories, such as active TB, QFT+, and QFT- individuals. Focusing on broader clinical TB states, ATB individuals showed elevated NK cell frequencies during treatment; however, consistent with previous reports (39,40), no significant differences were observed between ATB at diagnosis and QFT+/- groups. Intermediate monocytes, associated with inflammatory responses, were elevated in ATB individuals and declined during treatment, supporting their potential as markers of active disease. This aligns with previous findings showing increased intermediate monocytes in active TB compared to QFT+/- groups, while classical and non-classical monocyte populations remained unchanged (29). These findings suggest that intermediate monocytes may be sensitive indicators of active TB and treatment response. As expected, markers of recent activation, HLA-DR and CD38, were increased on both CD4+ and CD8+ T cells from individuals with active TB at diagnosis. Recent papers showed a multidimensional analysis of the activation state of TB-specific CD4+ T-cells reactive to the TB antigens PPD or ESAT-6/CFP-10 by expression of CD154, CD38, HLA-DR, and Ki-67 where they observed higher frequency of activated cells in TB disease compared to TB infection(41–43) The decline in these activation markers with treatment reflects a resolution of the immune activation associated with active infection. It is also supported by a recent study which showed a significant decline in CD38⁺HLA-DR⁺ PPD-specific CD4+ T cells after 8 weeks of anti-TB treatment, while the CD38⁻HLA-DR⁻ subset remained stable. These activated T cells were also highly expressed in tissue-resident memory populations at sites of active infection, suggesting that their decrease during treatment reflects reduced antigenic stimulation (42). However, these activated cells were similar in QFT subgroups, indicating no association of these activated cells with varying QFT values or exposure. Again, no differences were observed in NKT cells or T:M doublets in QFT subgroups, showing these subsets could only distinguish clinical states of TB. However, NKT cells showed an increase in ATB individuals after treatment, indicating a possibly distinct role for NKT cells during TB treatment. This increase likely reflects reduced antigenic burden and immune exhaustion as the infection is controlled. However, the extent and timing of this recovery can differ based on factors such as treatment duration, disease severity, and host immune status (e.g., HIV co-infection). A study had reported that NKT cell numbers and function are often reduced in active TB compared to healthy controls (44). In relation to this, T:M doublets were reduced in ATB individuals during treatment, suggesting their presence may be associated with active disease. A study by our group quantified the association constant (Ka) of T cell– monocyte complexes and reported a similar decline in Ka during treatment in active TB patients. These findings indicate that both the frequency and strength of T:M interactions may serve as potential biomarkers for monitoring treatment response and identifying individuals at risk of relapse (30). Together, these observations suggest that monitoring dynamic changes in immune cell interactions, particularly T:M and NKT cells, could offer valuable insight into TB disease activity and progression, but could not distinguish different QFT subgroups stratified based on longitudinal QFT values.

Another important aspect of the immune response against TB infection is the polarized phenotype of CD4 T cells. We observed differences in the frequency of Th subsets in clinical states of TB but observed comparable frequencies of these subsets in QFT subgroups. ATB individuals had lower frequencies of Th1 cells but higher frequencies of Th17 cells compared to QFT+ and QFT− controls, consistent with the pro-inflammatory role of Th17 cells in TB pathogenesis (45,46). Treatment was associated with a decline in Th1 and Th1* subsets and an increase in Th2 cells, reflecting a shift toward a more anti-inflammatory and tissue-repair-focused immune profile. In terms of chemokine expression, cells expressing CXCR3, irrespective of other chemokine receptors, declined with treatment, which may be associated with treatment and disease resolution. CXCR3+ T cells are being recruited in the lung where they can actively kill the bacteria and thus lead to disease resolution(47). Previously, our group has shown that Th1* (CCR6⁺, CXCR3⁺, CCR4⁻)cells exhibit a hybrid Th1/Th17 lineage signature along with a distinct transcriptional profile marked by genes linked to TB susceptibility, heightened T cell activation, increased survival, and cytotoxic functions resembling those of CTLs (8). These subsets also change depending on the tissue. A study demonstrated that Th1 cells are the dominant T helper subset in tuberculosis, appearing as low-differentiated CXCR3⁺CCR6⁺ cells in the blood and transitioning into highly differentiated CXCR3⁺/⁻CCR6⁻ cells in the lungs, highlighting their key role in TB immunity and the compartment-specific maturation of antigen-specific T cells (48). The altered Th1/Th2 ratio in ATB cases further underscores the immune dysregulation characteristic of TB (26,49). Similar to the above findings, no significant differences were observed across any of the analyzed Th subsets in QFT subgroups, indicating that variations in QFT levels do not correspond to shifts in Th cell distribution within this cohort. Overall, we observed notable differences in immune responses between active TB patients at the time of diagnosis and following treatment. However, such differences were not evident when participants were stratified into QFT subgroups based on the highest and lowest QFT values. Additionally, most immune cell subsets appeared comparable between the QFT+ and QFT- groups. This lack of significant distinction may be explained by the fact that the participants in both QFT groups were close contacts of TB patients, potentially resulting in shared antigen exposure and similar levels of immune priming or activation despite differences in QFT response. A recent study investigated the immunological characteristics of *M.tb* QuantiFERON-TB Gold (QFT) reverters, individuals who initially test QFT+ but later revert to QFT-. Compared to persistent QFT+ and QFT- individuals, reverters exhibited intermediate Mtb-specific Th1 responses. These included reduced proportions of stem cell memory and early differentiated IFN-γ⁻TNF⁺IL-2⁻ CD4⁺ T cells, suggesting partial immune control. However, no significant differences were observed in T cell activation marked by HLA DR expression(50). A positive QFT conversion during follow-up could suggest a higher risk of developing active TB. Based on this, we expected that these individuals might show immune changes similar to those seen in people with active TB, especially in markers of recent immune activation like HLA-DR⁺CD38⁺. We didn’t observe this pattern in our study, although, a larger group of participants would be needed to confirm these findings.

Our exploratory study has focused on immunological profiling, enabling the characterization of dynamic immune responses across the TB spectrum and treatment. However, the study has certain limitations. The small sample size in some groups, particularly ATB individuals at mid-treatment, may have limited the statistical power to detect subtle differences, and the lack of longitudinal measurements impairs the conclusions that can be drawn. The upregulation of activation markers following antigen-specific stimulation was measured, but the cytokine profile of the responding cells was not determined to further phenotype them. The frequencies of non-conventional T cells and whether they mediated a large proportion of the response against Mtb lysate were also not determined.

In conclusion, our findings provide insights into the immunological heterogeneity associated with Mtb infection and TB treatment. Importantly, our study showed that ex vivo phenotyping of close contacts of TB patients stratified based on longitudinal QFT values could not distinguish them in the spectrum of TB infection, or these individuals may not exhibit differences in their immune profile. Future studies should focus on validating these findings in larger, longitudinal cohorts and exploring the underlying mechanisms driving the immune responses.

## MATERIAL AND METHODS

### Study approval

All participants provided written informed consent for participation in the study. Ethical approval was obtained from the institutional review boards at Pneumology Institute (CE-3/2018) and the University of California San Diego (180068).

### Study participants

We conducted a prospective longitudinal cohort study to investigate the spectrum of TB infection(51). Recruitment occurred throughout the Republic of Moldova, with the exclusion of Transnistria, from October 1, 2018, through December 31, 2021. Individuals with newly diagnosed TB were designated as index cases and after recruitment were asked to identify close contacts to the best of their ability and recollection. These close contacts were monitored over a period of at least 24 months to determine rates of progression to TB disease. Screening for index cases was initiated through clinic microscopy centers and family doctors caring for referred patients and the national online TB registry, the System of Information for Monitoring and Evaluation of TB patients (SIME-TB), was also monitored for new TB diagnoses for potential recruitment. All close contacts were routinely reviewed for TB disease based on clinical evaluation by the participant’s primary treating physician, Moldova team physicians, and study physicians at the University of California, San Diego. The inclusion criteria for participants with pulmonary TB were (1) sputum acid-fast bacilli (AFB) smear-positive or nucleic acid amplification test (NAAT)-positive within the previous four weeks, (2) ≥18 years old, and (3) willing to identify contacts in the past three months, whether living with the contacts in the same house or not. TB patients were excluded from the study if they had received any TB treatment for more than 4 weeks prior to screening and 12 weeks prior to testing. The inclusion criteria for contacts were (1) exposure to a smear-positive or NAAT-positive TB patient, (2) ≥5 years old and ≥16 kg, (3) no evidence of TB on clinical evaluation which includes chest x-ray. Contacts were excluded if they (1) were pregnant, (2) were ever treated to prevent TB (TB preventative therapy is not routinely provided in the Republic of Moldova), (3) had a history of prior TB disease, (4) had their QFT-Plus tests after 12 weeks post exposure, or (5) had no QFT-Plus at baseline.

Active pulmonary tuberculosis (ATB) was diagnosed based on at least one of the following criteria: positive mycobacterial PCR result (GeneXpert MTB/RIF (Cepheid, Sunnyvale, CA, USA) or MGIT liquid culture. Close contacts with evidence of TB disease identified within 30 days of index case treatment initiation were considered “co-prevalent” cases given temporal proximity to the index case infectious period and were excluded from this analysis. Close contacts who had no evidence of clinical TB disease on recruitment and developed TB disease 30 days or more after index case treatment initiation were considered “progressor” cases. They were excluded from this study.

Each close contact was tested with a QFT plus at enrollment, and at 3, 6 and 12 months follow-up visits. The QFT was considered positive if the IFN-γ response to either Mtb antigen was significantly above the Nil value, using the recommended threshold of ≥0.35 IU/mL and ≥25% of the Nil value. Based on the QFT plus results, participants were divided into subsets to understand the spectrum of infection: QFT negative stable: individuals with QFT values below the cut-off value of 0.35 IU/mL and remained negative upon follow up. These are referred to as QFT- throughout the manuscript. Specifically, their maximum QFT plus value was less than 0.1, with little to no variation (less than 0.001), QFT Increasing: This group consists of QFT+ individuals whose QFT values increased over time. Their average QFT ranged between 0.35 and 1-1.5, with variation less than 0.5. QFT increasing individuals were also subdivided into QFT low (at baseline) and QFT high (at 6 months) (see figure 1). QFT Positive Stable: These individuals consistently maintained high QFT values over time. Their average QFT was greater than 1.5, with little variation (less than 0.5). Converters: These individuals converted from QFT- to QFT+ over time. All close contacts had no clinical signs and symptoms of TB throughout the 12 month period study. For this study, we initially had samples from 40 ATB individuals, 57 QFT increasing, 42 QFT stable, 41 converters, and 50 QFT- individuals. However, due to low cell viability, we restricted our analysis to samples with viability greater than 50%. The final cohort comprised 12 ATB individuals at diagnosis and 13 at mid-treatment. Among the QFT+ group, we selected 30 individuals from the QFT-increasing group—15 with the lowest baseline QFT values and 15 with the highest follow-up values (difference >0.5 IU/mL), 27 from the QFT-stable group, and 16 converters (8 at baseline with QFT- who later converted to QFT+). The QFT- included 16 individuals with QFT values consistently below 0.35 IU/mL.

### PBMC isolation and thawing

PBMCs were purified from whole blood by Ficoll gradient centrifugation and stored in Falcon (Becton, Dickinson and Company, Franklin Lakes, NJ, USA) and SepMate (STEMCELL Technologies Inc., Vancouver, Canada) tubes. Cells were resuspended in FBS (Gemini Bio-Products) containing 10% DMSO (v/v, Sigma) and cryopreserved in liquid nitrogen. Cryopreserved PBMC were quickly thawed by incubating each cryovial at 37°C for 2 min, and cells were transferred to complete RPMI medium which is RPMI 1640 with L-glutamin and 25 mM HEPES; Omega Scientific, supplemented with 5% human AB serum (GemCell), 1% penicillin streptomycin (Life Technologies), 1% glutamax (Life Technologies), and 20 U/ml benzonase nuclease (MilliporeSigma). Cells were centrifuged and resuspended in complete RPMI medium to determine cell concentration and viability using trypan blue. Samples with a viability >50% were used for downstream analysis, while the remaining samples were discarded.

### Ex vivo cell frequency by multicolor flow cytometry

After thawing and checking the viability of the PBMCs, 1x10^6^cells were plated in a 96-well plate. Cells were then washed with PBS to remove any remaining RPMI media. Subsequently, the cells were resuspended in 100μl solution consisting of 5μl Fc block (BD), 0.2ul Live/Dead dye eFluor506 (eBiosciences), and 94.8μl PBS and incubated for 20 minutes at room temperature. Following this incubation, the cells were washed with 100 μL of FACS buffer (PBS containing 10% FBS) and stained with an antibody cocktail comprising surface-expressed antibodies (details provided in supplementary table 2) for 20 minutes at room temperature. Post-incubation, the cells were washed twice with FACS buffer and finally resuspended in 100 μL of FACS buffer before being acquired on a Cytek Aurora spectral flow cytometer. Data analysis was performed using FlowJo version 10.1 (Treestar Inc.). Gating strategy is shown in Supplementary figure 1.

### Antigen-specific response by AIM assay

After thawing and checking the viability of the PBMCs, 1x10^6^ cells per condition were plated in a 96-well plate, and stimulated with MTB300 (2μg/ml) or MTB lysate (10ug/ml) for 24 hours in complete RPMI medium at 37°C with 5% CO_2_. PBMCs incubated with DMSO at the same concentration as in MTB300 were used as a negative control to assess non-specific background signal. Plate-bound anti-human CD3 antibody (clone OKT3, Invitrogen) and soluble anti-human CD28 antibody (clone CD28.2, BD Biosciences) at a final concentration of 1 μg/mL was used as a positive control. After 24 hours, cells were washed twice in FACS buffer and stained with surface-expressed antibodies (supplementary table 2) along with fixable live/dead stain for 20 -30 minutes at room temperature. After the incubation, cells were washed twice with FACS buffer and finally resuspended in 100 μL FACS buffer before acquiring on the Cytek Aurora spectral flow instrument. Data analysis was conducted in FlowJo version 10.1 (Treestar Inc.). The background-subtracted signal was calculated as the frequency of AIM+ cells in the antigen stimulation minus the frequency in the DMSO stimulated condition. Gating strategy is shown in Supplementary figure 2.

### Statistical analysis

Statistical analyses were performed using GraphPad Prism software (GraphPad Software, Inc., San Diego, CA, USA, version 9.2). Data is shown as median with interquartile range. A non-parametric test was applied after checking for normality. Median values of non-parametric data were compared between groups using Mann-Whitney tests (for comparing two groups), and Kruskal-Wallis tests (for comparing more than two groups), with Dunn’s post-test correction. P values are two-tailed, and a value of less than 0.05 was considered significant.

## Author contributions

C.S.L.A., B.P., J.B., A.C., T.R., and D.G.C. participated in the design and direction of the study. S.P., C.C., and J.B. performed and analyzed the experiments. V.C., N.C., N.H., and D.G.C. recruited participants, performed clinical evaluations, and isolated PBMCs. S.P., J.B., and C.S.L.A. wrote the manuscript. All authors read, edited, and approved the manuscript before submission.

